# The QxxR Motif of RNA Helicase Me31B Is Essential for Drosophila Female Fertility and Germline Development

**DOI:** 10.64898/2026.08.27.747641

**Authors:** Raheem Mansoor, Anum S Minhas, Abby N Thomas, Ammara A Mansoor, Aidan H McCambridge, Carol Dilts, John Eshak, Deep Govani, Brynn Nylin, Jonathan C Trinidad, Ayah Y Kanaan, Evan Kara, Abraham Fielder, Isaac Fielder, Adriana Iglendza, Yousif Mukatash, Brooke Pumnea, Melissa M Menzel, Ahad L Shabazz Henry, Matthew G Niepielko, Ming Gao

**Author notes:** Correspondence: M. Gao., M. G. Niepielko.

## Abstract

The QxxR motif is evolutionarily conserved within DEAD-box RNA helicases, including *Drosophila* Me31B and human DDX6, which post-transcriptionally regulate gene expression during animal development. A pathogenic H372R substitution (Qx<u>H</u>R to Qx<u>R</u>R) in the QxxR motif of human DDX6 has been associated with various developmental defects, but how this motif contributes to DDX6-family protein function remains unclear. Here, we used *Drosophila* Me31B as an *in vivo* model to investigate the QxxR motif’s developmental role. We generated a *Drosophila* strain carrying the corresponding H333R missense mutation in Me31B and characterized its effects on female fertility, embryonic viability, germline development, and Me31B-associated molecular pathways. The *me31B^H333R^* mutation reduced female fertility in a gene dose-dependent manner, with homozygous mutant females being sterile. Embryos from the mutant females also exhibited primordial germ cell defects. Despite these developmental phenotypes, the *me31B^H333R^* mutation did not significantly alter Me31B protein abundance, global ovarian transcriptome or proteome profiles, or representative germ plasm mRNA and protein localization. In contrast, bait-normalized IP-MS analysis revealed altered enrichment of selected Me31B-associated proteins, including increased association of known Me31B interactors Trailer hitch (Tral) and Ypsilon Schachtel (Yps). These findings establish Me31B^H333R^ as an *in vivo* model for investigating the conserved QxxR motif and suggest that disruption of this motif compromises development not through broad changes in gene expression, but potentially through altered composition or regulation of Me31B-containing ribonucleoprotein complexes.

## Introduction

Me31B is the *Drosophila* ortholog of the conserved DDX6 family of DEAD-box RNA helicases, a protein family with central roles in post-transcriptional gene regulation [1–9]. DDX6-family proteins are conserved across metazoans and regulate diverse aspects of post-transcriptional gene expression, including mRNA storage, transport, translational repression, stabilization, and decay (reviewed in [10, 11]). Through these coordinated activities, they ensure that maternal and developmental transcripts are expressed at the precise time and location required for proper development [6, 11–14].

Like other DEAD-box helicases, Me31B contains conserved helicase-core motifs that contribute to RNA and ATP binding, ATP hydrolysis, and conformational changes associated with RNA remodeling, and several of these motifs are required for Me31B function *in vivo* [15–17]. Although the functions of several canonical helicase-core motifs have been studied in detail, the biological role of the conserved QxxR motif remains poorly understood. Previous structural and biochemical analyses suggest that this motif contributes to RNA interaction, implying a potential role in RNA substrate engagement or helicase regulation [18], but its developmental function *in vivo* remains unknown.

The biological importance of this motif is further supported by human genetic evidence. A de novo missense mutation in human DDX6, H372R, which converts the QE**<u>H</u>**R sequence to QE**<u>R</u>**R within the conserved QxxR motif, was identified in an individual with intellectual disability, dysmorphic features, genitourinary anomalies, and developmental delay [19]. This finding suggests that the QxxR motif is functionally important for DDX6 activity *in vivo*. However, because DDX6 acts in many tissues and regulatory contexts, the mechanistic basis by which this motif supports development remains difficult to resolve in humans alone.

Here, we use *Drosophila* Me31B as an *in vivo* model to investigate the developmental function of the conserved QxxR motif. The female germline provides a particularly relevant system because oogenesis and early embryogenesis depend on post-transcriptional regulation of maternally supplied RNAs, which must be stored, localized, repressed, or translated in spatially and temporally controlled patterns [20, 21].

Consistent with this role, DDX6-family proteins are enriched in germline ribonucleoprotein (RNP) particles and are required for fertility and early developmental programs in multiple organisms [22–24]. In *Drosophila*, Me31B is an abundant component of maternal mRNP complexes and has established roles in oogenesis, maternal mRNA regulation, and early embryonic development [23, 25–27]. Therefore, we generated an H333R mutation in the Me31B QxxR motif, converting Qx**<u>H</u>**R to Qx**<u>R</u>**R and thereby reproducing the human DDX6 H372R substitution, to determine how disruption of this motif affects germline development and Me31B-associated regulatory pathways.

Here, we show that the *me31B^H333R^* mutation causes severe female fertility defects, embryonic hatchability failure, and germline development defects. Despite these severe developmental phenotypes, the mutation produces only limited changes in germ plasm organization and global ovarian RNA and protein abundance. Instead, our data suggest that the H333R mutation alters the composition or regulation of Me31B-containing RNP complexes. Together, these findings establish a *Drosophila* model for investigating the conserved QxxR motif of DDX6-family RNA helicases, providing insights into how its mutation disrupts Me31B function *in vivo*.

## Results

### Mutation in the Me31B QxxR motif decreases *Drosophila* female fertility

To investigate the *in vivo* function of the conserved QxxR motif of Me31B, we used CRISPR-mediated genome editing to generate a H333R missense mutation within the QxxR motif (Qx**<u>H</u>**R → Qx**<u>R</u>**R). This substitution corresponds to the human DDX6 H372R mutation (Supplementary Figure S1), which is associated with developmental defects, allowing us to model QxxR motif disruption in *Drosophila*. Given the established role of Me31B in female germline development [10, 15], we first tested whether the *me31B^H333R^* mutation affects female fertility.

We performed a fertility assay by crossing individual *me31B^H333R^* mutant females to wild-type *w^1118^* males and quantified the number of eggs laid, viable progeny produced, and egg hatch rate (Supplementary Table S1). Both homozygous and heterozygous *me31B^H333R^* females were analyzed to determine whether the phenotype(s) depended on mutant-allele copy number, using genotype-matched *me31B^WT^* females as controls. The *me31B^WT^* control strain was generated using the same CRISPR strategy without altering the Me31B coding sequence and was previously shown to display wild-type-like fertility [15].

The assays showed that homozygous *me31B^H333R^* females laid an average of 37 eggs over the 10-day assay period, compared to 92 eggs laid by homozygous *me31B^WT^* control females, corresponding to a 60% reduction that did not reach statistical significance (Figure 1A). Heterozygous *me31B^H333R^* females laid an average of 48 eggs, compared with 78 eggs laid by heterozygous *me31B^WT^* controls, corresponding to a 39% reduction that was also not statistically significant (Figure 1A). In contrast, viable progeny production was strongly reduced. Homozygous *me31B^H333R^* females produced an average of only 1 progeny, compared to 81 progeny from homozygous *me31B^WT^* controls, while heterozygous *me31B^H333R^* females produced an average of 20 progeny, compared with 71 progeny from heterozygous *me31B^WT^* controls (Figure 1B). Both reductions were statistically significant, although the difference in progeny number between homozygous and heterozygous *me31B^H333R^* females was not.

**Figure 1.**
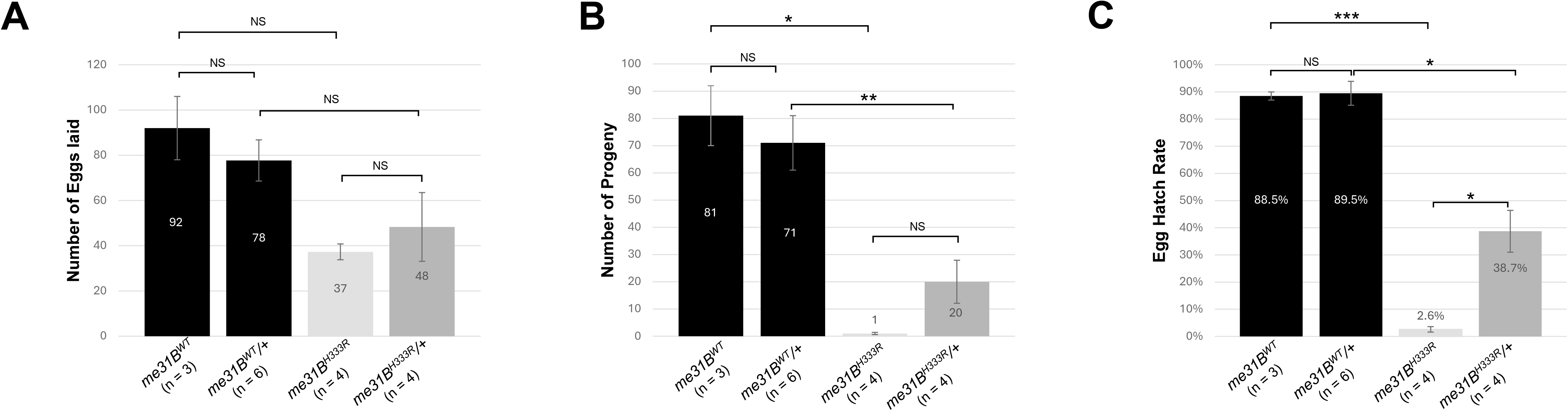
The fertility assay performed with *me31B^H333R^* and *me31B^WT^* control females. **(A)** Quantification of the number of eggs laid by a female fly from strains *me31B^H333R^*, *me31B^H333R^*/+, *me31B^WT^* (homozygous control), and *me31B^WT^*/+ (heterozygous control). **(B)** Quantification of the number of viable progeny produced by the female flies from strains *me31B^H333R^*, *me31B^H333R^*/+, *me31B^WT^*, and *me31B^WT^*/+. **(C)** Quantification of egg hatch rate in female flies from strain *me31B^H333R^*, *me31B^H333R^*/+, *me31B^WT^*, and *me31B^WT^*/+. Error bars represent the standard error of the mean. **n r**epresents the number of biological replicates (full data in Supplementary Table S1). **NS**, not significant. **\***, **\*\***, and **\*\*\*** represent p-values less than 0.05, 0.01, and 0.001, respectively.

We further calculated egg hatch rates (see Materials and Methods) and found that they were significantly decreased in mutant females. Eggs from homozygous *me31B^H333R^* females showed a hatch rate of 2.6%, compared with 88.5% for homozygous *me31B^WT^* controls, while eggs from heterozygous *me31B^H333R^* females showed a hatch rate of 38.7%, compared with 89.5% for heterozygous *me31B^WT^* controls (Figure 1C). The hatch rate was also significantly lower in homozygous *me31B^H333R^* females than in heterozygous *me31B^H333R^* females (Figure 1C), indicating that the mutation impairs egg viability in a gene dose-dependent manner.

Together, these results show that the Me31B QxxR motif mutation reduces *Drosophila* female fertility primarily by decreasing the hatchability of eggs produced by mutant females. The significant fertility defect in heterozygous females, along with the more severe phenotype in homozygous females, indicates that the *me31B^H333R^* mutation acts as a dose-sensitive allele for female fertility, particularly egg hatchability.

### Homozygous *me31B^H333R^* females are sterile due to egg hatchability failure

The near-complete loss of progeny production in homozygous *me31B^H333R^* females in the initial fertility assay suggested that the one rare viable progeny observed could represent an exceptional event rather than typical reproductive output. To test whether homozygous *me31B^H333R^* females are effectively sterile, we performed a grape agar-based egg hatchability assay with more embryos and directly monitored egg development.

In this assay, none of the 250 eggs laid by homozygous *me31B^H333R^* females developed into larvae or later stages, corresponding to a 0% hatch rate (Figure 2A). In contrast, 91.8% of the 550 eggs laid by *me31B^WT^* control females hatched successfully (Figure 2A). Despite this complete hatchability failure, eggs laid by homozygous *me31B^H333R^* females appeared externally similar to control eggs under dissection microscopy (Figure 2B). These observations indicate that sterility in homozygous *me31B^H333R^* females is primarily associated with embryonic developmental failure rather than obvious defects in external egg morphology. These data demonstrate that homozygous disruption of the Me31B QxxR motif causes female sterility, with mutant females producing eggs that appear morphologically normal but fail to hatch.

**Figure 2.**
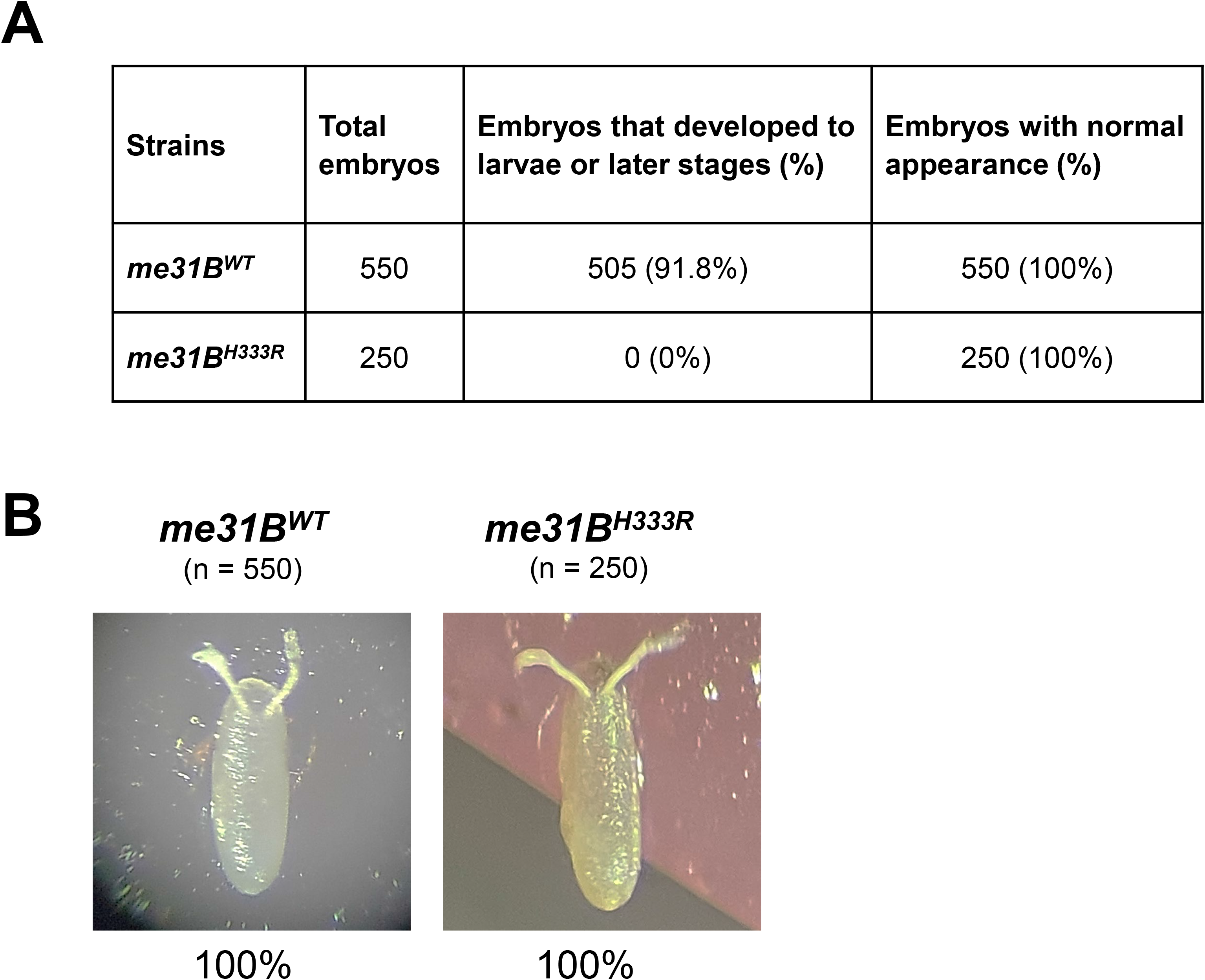
Hatchability assay and external morphology of eggs laid by *me31B^H333R^* and *me31B^WT^* control females. **(A)** A total of 550 eggs from *me31B^WT^* control females and 250 eggs from *me31B^H333R^* mutant females were analyzed in the grape agar-based hatchability assay. None (0%) of the eggs from the *me31B^H333R^* mutant developed to larvae or later stages, while 505 (91.8%) of those from the control developed to larvae or later stages. Egg appearance from both strains was examined and shown in (B). **(B)** Representative images of eggs laid by *me31B^WT^* control females and *me31B^H333R^* mutant females. Eggs from the *me31B^H333R^* mutant females look similar in appearance to those of the control. **n** represents the number of eggs analyzed from each strain.

### Defective primordial germ cell phenotypes in *me31B^H333R^* Mutants

We next investigated whether the *me31B^H333R^* mutation affected germline development by monitoring primordial germ cells (PGCs) in the embryos derived from *me31B^H333R^* mothers. Using anti-Vasa immunofluorescence to label PGCs and confocal microscopy, we observed an increase in defective PGCs that failed to migrate and coalesce into the embryonic gonads in both homozygous and heterozygous mutants (Figure 3).

**Figure 3.**
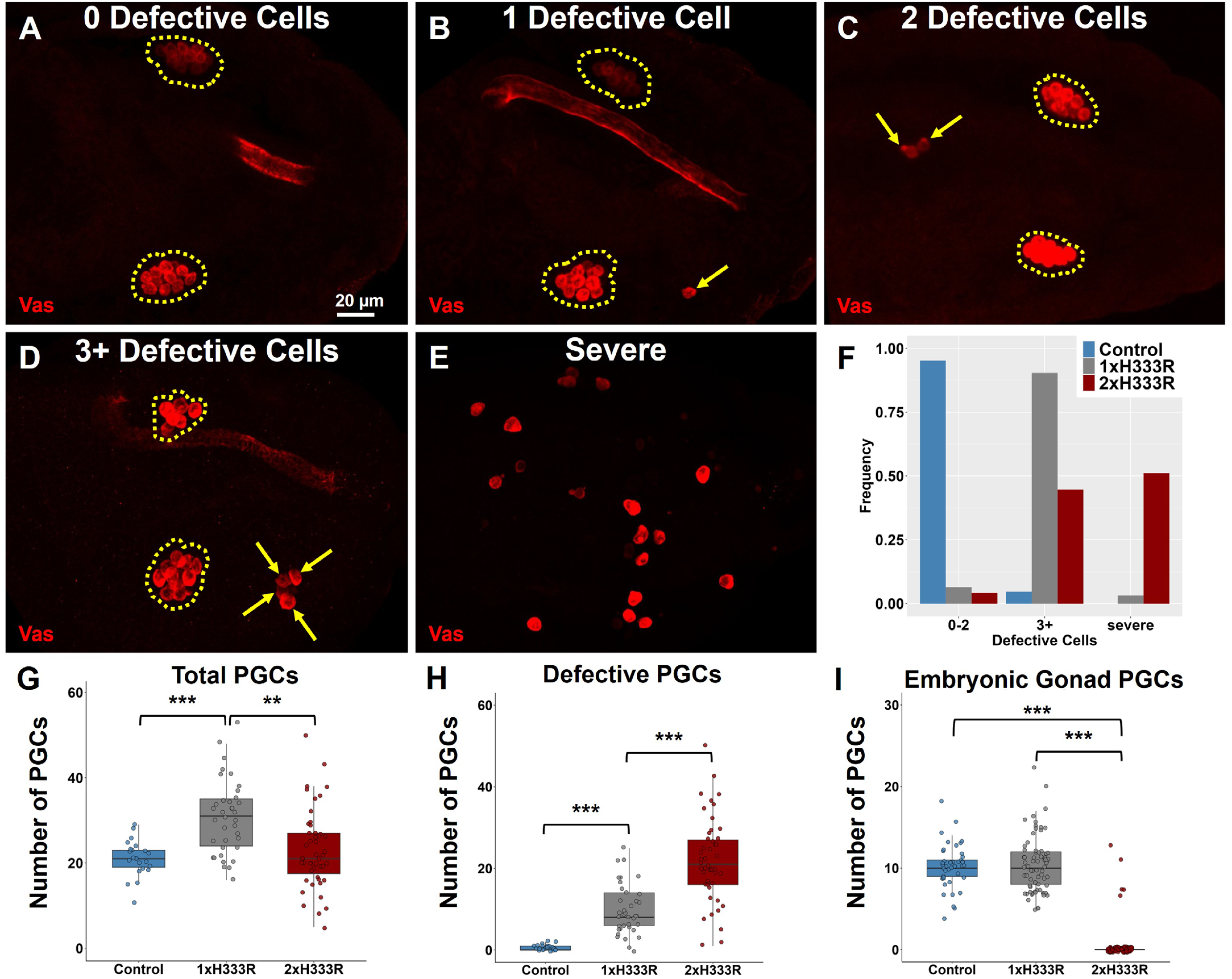
Visualization and quantification of defective primordial germ cell migration in control and mutant *me31B* embryos. **(A-D)** Representative 40x confocal microscopy images of embryos with (A) no defective PGCs, (B) one defective PGC, (C) two defective PGCs, and (D) three or more defective PGCs, respectively. Embryonic gonads are outlined by yellow dotted circles. Defective PGCs that have not properly migrated to the gonads are indicated with yellow arrows. **(E)** Representative 40x confocal microscopy image of an embryo with a severe phenotype characterized by the absence of properly formed embryonic gonads and scattered PGCs. **(F)** Histogram showing frequency of embryos with 0-2 defective PGCs, 3+ defective PGCs, or severe phenotype in control, *me31B^H333R^*/+, and *me31B^H333R^* flies. n = 20 in control, n = 47 in *me31B^H333R^*, and n = 23 in *me31B^H333R^*/+ embryos. Control, 1x H333R, and 2x H333R represent genotypes *me31B^WT^*, *me31B^H333R^*/+, and *me31B^H333R^*, respectively. **(G-I)** Primordial germ cell (Vasa-positive cells) counts for Control, 1x H333R, and 2x H333R genotypes, where (G) represents total PGC counts, (H) shows counts for defective PGCs (Vasa-positive cells outside of the embryonic gonad), and (I) shows the number of PGCs that have coalesced within an embryonic gonad, n > 22 embryos. Statistical significance was determined using an ANOVA test with a Tukey post-hoc test in R using the DescTools package.

Specifically, 95.2% of *me31B^WT^* control flies had either none or one to two defective germ cells, with only 4.8% of embryos having three or more defective cells (Figure 3A-D; 3F). Furthermore, none of the control embryos had a severe-defective phenotype (Figure 3E-3F). In contrast, the *me31B^H333R^*/*+* mutant flies displayed a significant increase in the number of defective cells seen within the embryos of these flies: 6.5% of the embryos observed had 0-2 defective cells (Figure 3A-C; 3F), while 90.3% of the embryos had three or more defective cells (Figure 3D; 3F) and 3.2% of the embryos had a severe-defective phenotype (Figure 3E-3F). In *me31B^H333R^* mutant flies, very few embryos (4.3%) had zero to two defective cells (Figure 3A-C; 3F), while 51% had a severe-defective phenotype, and the rest (44.7%) had three or more defective cells (Figure 3D-3F). Altogether, these data demonstrate that the *me31B^H333R^* mutation affects germline development. Furthermore, the increased frequency of severely defective germ cells from heterozygous to homozygous mutants suggests that the *me31B^H333R^* mutation has a gene-copy dosage effect.

Next, we investigated how these genotypes affected 1) the total number of PGCs, 2) the total number of defective PGCs, and 3) the number of PGCs within an embryonic gonad in 18-24hr embryos (Figure 3 G-I). We found an average of 21.1 ± 0.9, 30.7 ± 1.4, and 23.1 ± 1.4 total PGCs in the control, *me31B^H333R^*/+, and *me31B^H333R^* mutant, respectively (Figure 3G). The increase in total PGCs in the *me31B^H333R^*/+ mutant at this stage was significant when compared to the control (*p* < 0.001), and when compared to the *me31B^H333R^* mutant (*p* < 0.002). No change was observed between the control and the *me31B^H333R^* mutant (*p* = 0.47). On average, we found 0.59 ± 0.16, 10.0 ± 1.0, and 22.2 ± 1.5 total defective PGCs in the control, *me31B^H333R^*/+, and *me31B^H333R^* mutant, respectively (Figure 3H). Compared to the control, the increase in total number of defective PGCs in the *me31B^H333R^*/+ and *me31B^H333R^* mutants at this stage was significant (*p* < 0.001), as was the difference between the *me31B^H333R^*/+ and *me31B^H333R^* mutants (*p* < 0.001). Next, we analyzed how the H333R mutation affected the average number of PGCs found within an embryonic gonad. We found averages of 10.25 ± 0.40, 10.4 ± 0.40, and 0.50 ± 0.22, in the control, *me31B^H333R^*/+, and *me31B^H333R^* mutant, respectively (Figure 3I). No difference was observed between the control and the *me31B^H333R^*/+ mutant (*p* = 0.92), and these averages agree with what is observed in wild-type *D. melanogaster* [28]. However, the lack of PGCs in the embryonic gonad of the *me31B^H333R^* mutant was significant compared with the control (*p* < 0.001). Altogether, we found that the *me31B^H333R^*/+ mutant has the propensity to increase the PGCs without increasing the numbers within the embryonic gonad, which is reflected in an increased presence of defective PGCs. In the *me31B^H333R^* mutant, our statistical analysis supports a model where increasing the H333R allele copy from one to two increases the number of defective PGCs by hindering their ability to coalesce into the embryonic gonad (see Discussion).

### Absence of PGCs observed in *me31B^H333R^* mutants

While viewing images of Vasa-labeled PGCs, we observed that some *me31B^H333R^* mutants’ embryos lacked PGCs altogether. Specifically, the *me31B^WT^* control flies had 97.4% ± 1.06% embryos with PGCs present (Figure 4A; 4D). In comparison, 79.5% ± 3.31% of *me31B^H333R^*/+ mutants had embryos with PGCs present, and *me31B^H333R^* mutants only had 38.9% ± 5.18% of their embryos with PGCs present (Figure 4B-D). Next, we aimed to determine whether the lack of PGCs was a direct effect on germline development or a more global effect on embryogenesis by co-staining with DAPI to mark somatic embryo nuclei and capture the progression of embryonic development (Figure 4E-G’). In all cases, embryos lacking PGCs also lacked DAPI in somatic embryonic cells; when PGCs were present, DAPI was also visible in embryonic somatic cells along with the presence of later embryonic development, regardless of whether the PGCs were coalesced within the embryonic gonad or had a severe PGC phenotype (Figure 4E-G’). These data indicate that when PGCs were not developing in *me31B^H333R^* mutants, the embryos were likely dead prior to fixation, suggesting that the absence of PGCs is likely due to the embryo not developing in general. These data are consistent with our earlier data that many mutant embryos failed to hatch and suggest that the failure likely resulted from early developmental defects.

**Figure 4.**
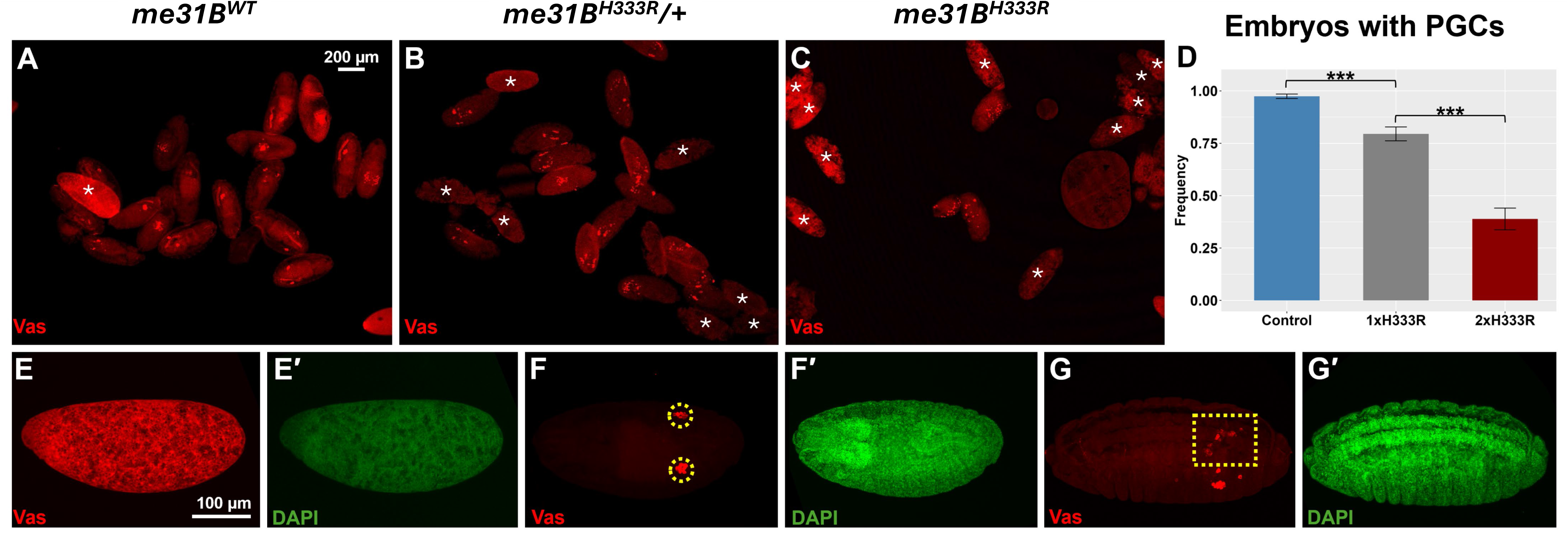
Loss of primordial germ cells in control and mutant *me31B* embryos. **(A)** Confocal microscopy image (4x) of Vasa-labeled PGCs in late-stage embryos of *me31B^WT^* flies. **(B)** Confocal microscopy image (4x) of Vasa-labeled PGCs in late-stage embryos of *me31B^H333R^*/+ flies. **(C)** Confocal microscopy image (4x) of Vasa-labeled PGCs in late-stage embryos of *me31B^H333R^* flies. Embryos lacking primordial germ cells are marked with an asterisk. **(D)** Bar graph showing frequency of embryos with PGCs present in control, *me31B^H333R^*/+, and *me31B^H333R^* flies. To calculate D, we determined the fraction of Vasa-positive embryos within a single low-magnification image and averaged it from >10 images gathered from 3 replicated experiments. Statistical significance was determined using an ANOVA test with a Tukey post-hoc test in R programming using the DescTools package, *p* < 0.005. **(E-E’)** Embryos stained with Vasa and DAPI show that embryos lacking PGCs are also DAPI negative for other embryonic cells. **(F-F’)** PGCs that have properly migrated to the embryonic gonads (yellow dotted circles) are DAPI positive. **(G-G’)** In embryos with defective PGCs (rectangle), DAPI staining is positive. Control, 1x H333R, and 2x H333R represent genotypes *me31B^WT^*, *me31B^H333R^*/+, and *me31B^H333R^*, respectively.

### The *me31B^H333R^* mutation does not significantly alter Me31B protein abundance

Because Me31B functions as a maternal regulator whose protein is produced during oogenesis and deposited into developing eggs [23, 29], we next asked whether the H333R mutation affects Me31B protein abundance in the ovary, which could simply explain for the observed fertility, hatchability, and PGC phenotypes. Since both the *me31B^H333R^* mutant and *me31B^WT^* control alleles encode C-terminally GFP-tagged Me31B proteins, we quantified Me31B levels in ovary lysates by anti-GFP Western blotting. Me31B^H333R^-GFP abundance was 118% ± 2.2% of the Me31B^WT^-GFP control level after normalization to the tubulin loading control, and this difference was not statistically significant (Figure 5). These results indicate that the H333R mutation does not significantly reduce Me31B protein abundance during oogenesis. Thus, the fertility, hatchability, and primordial germ cell phenotypes associated with *me31B^H333R^* are unlikely to be caused by a major loss of Me31B protein expression.

**Figure 5.**
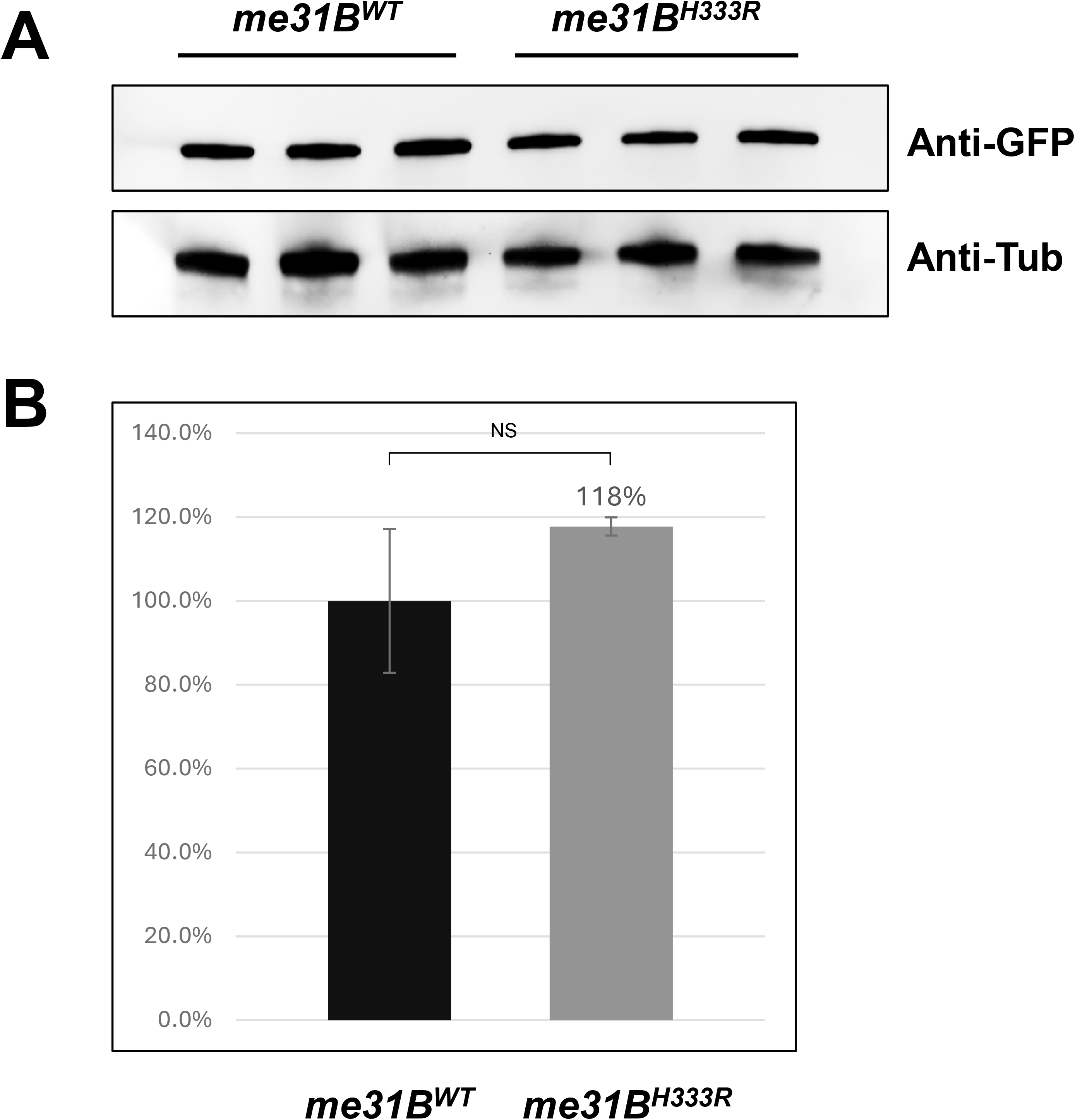
Western blot quantification of Me31B protein levels in *me31B^H333R^* and *me31B^WT^* control strains. **(A)** Anti-GFP Western blots were conducted to quantify the Me31B^H333R^-GFP and Me31B^WT^-GFP proteins from the ovaries of the corresponding strains. Anti-tubulin Western blots were used as loading controls. Three biological replicates are shown. **(B)** The expression levels of Me31B^H333R^-GFP proteins are 117.8% of those of the Me31B^WT^-GFP control proteins but are not significantly different. Uncropped images are provided as Supplementary Figure S2. Image analysis was performed with ImageJ, and protein quantification was normalized using alpha-tubulin. NS, not statistically significant. Error bars represent the standard error of the mean.

### Germ plasm localization of *nos* and *pgc* mRNA and Nos protein are unaffected in

### me31B^H333R^ mutants

Germline and embryonic development depend on the proper assembly of germ granules, as well as the proper translational regulation of their mRNA components, in the posterior germ plasm of the oocyte during embryogenesis [30]. Germ plasm formation requires the formation of thousands of ribonucleoproteins called germ granules, which contain homotypic clusters of different germ plasm mRNAs. Homotypic clusters are organized accumulations of identical mRNA transcripts that self-organize into discrete regions of a germ granule [31–33]. Given our previous research demonstrating that certain *me31B* mutations can affect the germ plasm localization of *nos* by reducing its homotypic cluster sizes within germ granules [15], we next performed smFISH to visualize and quantify potential effects on the germ plasm mRNAs *nanos* (*nos)* and *polar granule component* (*pgc*) in *yw* (control), *me31B^WT^* (control), *me31B^H333R^*/*+*, and *me31B^H333R^* embryos. In all four fly strains, *nos* and *pgc* mRNA transcripts were localized to the germ plasm and formed homotypic clusters. Using the Germ Granule Census (see Materials and Methods), we determined the average *nos* and *pgc* homotypic cluster content found in each germ granule. For *nos* mRNA, the average cluster size was 11.39 for *yw*, 11.18 for *me31B^WT^*, 11.36 for *me31B^H333R^*/+, and 11.39 for *me31B^H333R^* (Figure 6A-D’’). For *pgc* mRNA, the average cluster size was 6.63 for *yw*, 6.29 for *me31B^WT^*, 7.37 for *me31B^H333R^*/+, and 6.97 for *me31B^H333R^* (Figure 6A-D’’). Overall, we observed no significant differences in wild-type versus mutant mRNA localization or germ granule content, suggesting that the *me31B^H333R^* mutation causes developmental defects independent of germ plasm development and RNA localization.

**Figure 6.**
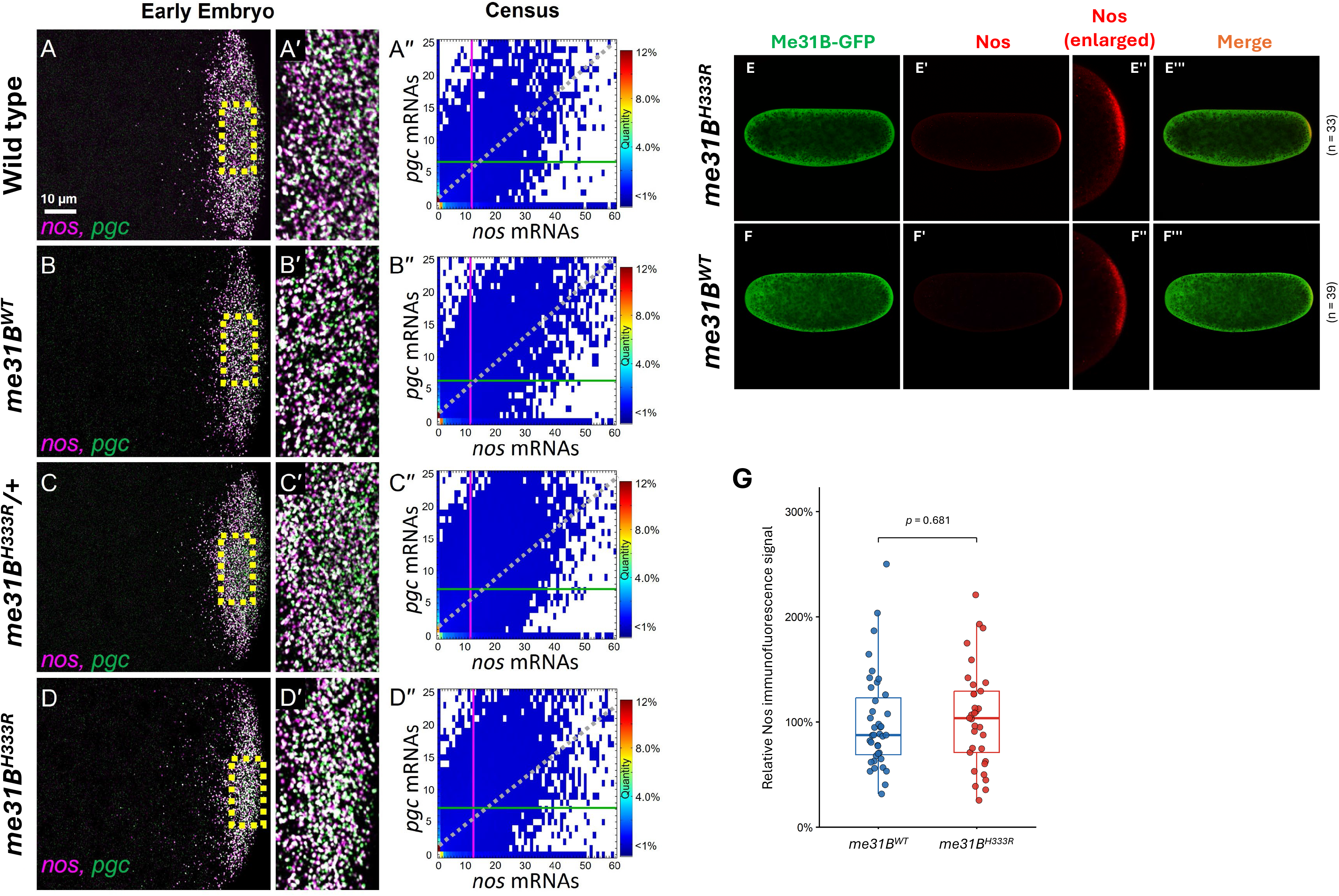
smFISH analysis of *nos* and *pgc* mRNA in wild-type, *me31B^WT^*, *me31B^H333R^*/+, and *me31B^H333R^* embryos and microscopic analysis of Nos protein in *me31B^H333R^* mutant and *me31B^WT^* control embryos. **(A-D)** Confocal images of early embryos of wild-type, *me31B^WT^*, *me31B^H333R^*/+, and *me31B^H333R^* flies showing localized *nos* (magenta) and *pgc* (green) mRNA transcripts in the posterior germ plasm. (A’-D’) Enlarged image of *nos* and *pgc* homotypic clusters highlighted in the yellow dotted boxes in the first column. (A’’-D’’) Germ Granule Census analyses of wild type, *me31B^WT^*, *me31B^H333R^*/+, and *me31B^H333R^ nos* and *pgc* germ granule content. The vertical magenta line represents the average number of *nos* transcripts within a germ granule. The green horizontal line represents the average number of *pgc* transcripts. The grey dotted line represents the balance of transcript numbers within co-localized *nos* and *pgc* homotypic clusters. Census data represents a pooled collection of n > 10,000 clusters across ≥ 6 different embryos. Significance was determined by calculating p-values comparing the average cluster sizes from ≥ 6 embryo replicates, which were > 0.90 and > 0.25 for *nos* and *pgc* cluster sizes, respectively, when compared to *yw* using an ANOVA test with a Dunnett’s post-hoc test using R statistical programming within the DescTools package. **(E-F)** Representative *me31B^H333R^* embryo (E -E’’’) and *me31B^WT^* control (F – F’’’) are visualized by endogenous GFP tag on Me31B proteins (green, E and F) and anti-Nos immunostaining (red, E’ and F’; E’’ and F’’ are enlarged images of posteriorly localized Nos in E’ and F’, respectively). Merged green and red channels are shown in E’’’ and F’’’. The posterior localization of Nos protein appear similar between the mutant and the control (compare E’ to F” or E’’ to F’’). **(G)** Quantification of Nos immunofluorescence signal within the posterior germ plasm. Each point represents one embryo (*me31B^WT^*, *n* = 39; *me31B^H333R^*, *n* = 33), and values were normalized to the mean of the *me31B^WT^* control, defined as 100%. Boxes indicate the median and interquartile range; whiskers represent the Tukey 1.5 × IQR range. The two genotypes did not differ significantly (two-sided Welch’s *t*-test, *p* = 0.681).

Given that Me31B has established roles in post-transcriptional regulation of maternal mRNAs, we asked whether the *me31B^H333R^* mutation disrupts the regulation of a representative germline determinant, Nos. Nos is a key germline protein whose spatially restricted accumulation in the early embryonic germ plasm depends on regulated expression of maternally supplied *nos* mRNA [34]. Previous studies have implicated Me31B-containing mRNP complexes in the repression and regulation of *nos* mRNA [35], making Nos a relevant readout for testing whether the *me31B^H333R^* mutation causes any major defect in this regulatory pathway.

We examined Nos protein expression and localization by immunostaining early embryos from *me31B^H333R^* mutant and *me31B^WT^* control females. This allowed us to assess Nos spatial distribution and relative signal patterns in the embryo germ plasm.

Immunostaining revealed posterior enrichment of Nos protein in the embryo germ plasm of *me31B^H333R^* mutant, with a localization pattern comparable to the control (Figure 6E-F’’’). Quantification showed that the mean Nos fluorescence signal of mutant embryos was 104.6% of the control and did not differ significantly between genotypes (*p* = 0.681; Figure 6G). Thus, the *me31B^H333R^* mutation does not cause an obvious disruption of Nos protein posterior localization in early embryos. These observations suggest that the reduced hatchability and germline phenotypes associated with the *me31B^H333R^* mutation are unlikely to result from a gross defect in Nos protein expression or localization.

Because Nos represents a single translationally regulated target and immunostaining provides an endpoint readout of protein accumulation rather than a direct measurement of translation, these data do not exclude more subtle effects on *nos* mRNA regulation or effects on other Me31B-regulated transcripts. Nevertheless, together with the limited proteome-wide perturbation detected in whole-ovary proteomics (next section), the preserved Nos pattern supports the interpretation that *me31B^H333R^* does not cause a broad collapse of this representative developmental control pathway.

### The *me31B^H333R^* mutation causes limited changes in the ovarian transcriptome and proteome

To search for molecular changes that could underlie the fertility, hatchability, and germline development phenotypes caused by the *me31B^H333R^* mutation, we next asked whether the mutation broadly alters gene expression in the female germline. We focused on ovaries because Me31B is produced during oogenesis and functions in germline RNP complexes involved in post-transcriptional regulation of maternal RNAs [23, 29, 36].

We performed whole-ovary RNA sequencing on ovaries from homozygous *me31B^H333R^* mutant and *me31B^WT^* control strains. The complete differential expression dataset is provided in Supplementary Table S2. Principal Component Analysis (PCA) of expression values showed separation between mutant and control samples along the first principal component, with some within-genotype variation among biological replicates (Figure 7A). Differential expression analysis identified 270 genes that differed significantly between *me31B^H333R^* and *me31B^WT^* control when applying both a statistical threshold (adjusted *p* < 0.05) and a fold-change cutoff (|log2 fold change| > 1), as visualized by a volcano plot (Figure 7B). Among these differentially expressed genes in the mutant, 159 were upregulated, and 111 were downregulated. These changes represent a relatively small fraction of the total 8,113 detected transcripts, while most transcripts showed comparable steady-state abundance between genotypes (Figure 7B).

**Figure 7.**
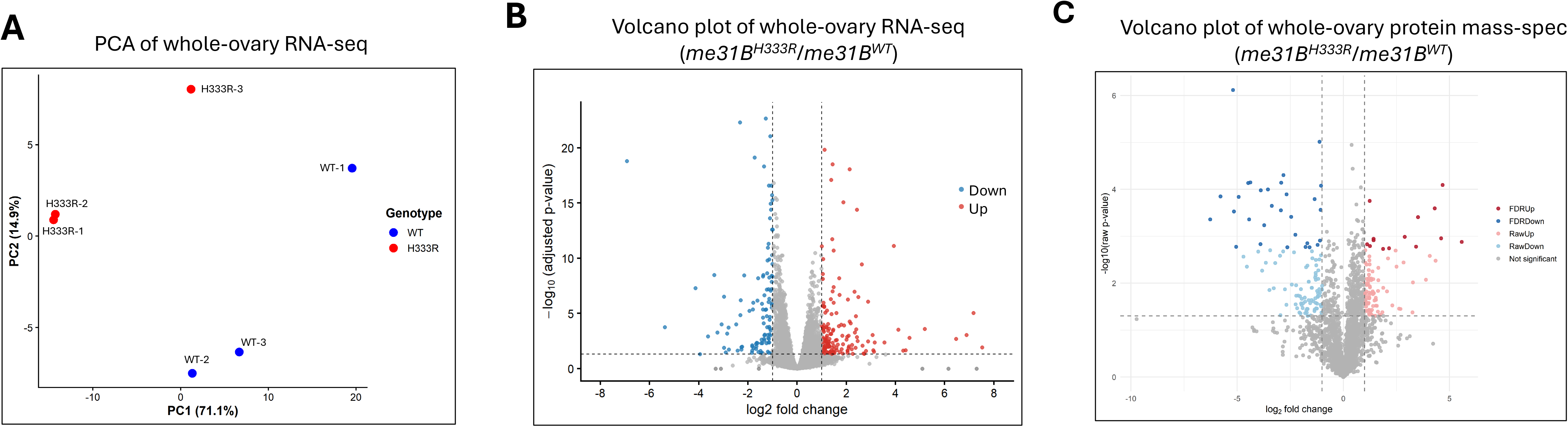
Whole-ovary RNA sequencing and proteomic analysis showing differential gene expression between *me31B^H333R^* and *me31B^WT^* control ovaries. **(A)** PCA of ovarian RNA-seq samples from *me31B^H333R^* and *me31B^WT^* control females. Each point represents one biological replicate, with three biological replicates per genotype. Percent variance explained by PC1 and PC2 is indicated on the axes. **(B)** Volcano plot showing differential gene expression between *me31B^H333R^* and *me31B^WT^* control ovaries. The x-axis indicates log2 fold change (*me31B^H333R^*/*me31B^WT^*) and the y-axis indicates −log10 (adjusted p-value). Genes meeting significance criteria (adjusted *p* < 0.05 and |log2 fold change| > 1) are highlighted in red (Up) or blue (Down), while all other genes are shown in grey. Dashed lines indicate fold-change and significance thresholds. **(C)** Volcano plot of differential protein abundance in whole-ovary lysates. Proteins detected by mass spectrometry in *me31B^H333R^* mutant and *me31B^WT^* control ovaries are plotted according to log2 fold change (*me31B^H333R^*/*me31B^WT^*) and −log10 (raw p-value). Proteins meeting stringent significance criteria (FDR < 0.05 and |log2 fold change| ≥ 1) are shown in dark red or dark blue. Proteins meeting raw p-value significance only (raw p < 0.05 and |log2 fold change| ≥ 1) are shown in light red or light blue. Grey points represent proteins that did not meet significance thresholds. Dashed lines indicate raw *p* = 0.05 and |log2 fold change| = 1.

To assess whether the differentially expressed mRNAs were enriched for specific functional categories, we performed Gene Ontology (GO) enrichment analysis using all genes detected in the RNA-seq dataset as the background. Neither the upregulated nor the downregulated gene sets showed significant enrichment for Biological Process or Molecular Function terms. These results indicate that the transcriptome changes associated with the *me31B^H333R^* mutation are limited and do not define a strongly enriched functional category at the level of steady-state RNA abundance.

To address whether the *me31B^H333R^* mutation causes broad changes in the ovarian proteome, we performed whole-ovary mass spectrometry and observed a small number of proteins with large abundance changes, whereas most detected proteins showed minimal differences between genotypes (Figure 7C; Supplementary Table S3). Specifically, of the 2,077 detected proteins, 44 met stringent criteria for differential abundance, defined by FDR < 0.05 and |log2 fold change| ≥ 1. GO enrichment analysis of the 44 differentially abundant proteins did not identify any significantly enriched functional categories.

Finally, comparison between the RNA-seq and proteomics datasets revealed minimal overlap between differentially expressed transcripts and differentially abundant proteins. Three genes showed concordant changes at both the transcript and protein levels: *TotA* and *TotC* (two members of the stress-inducible *Turandot* gene family [37]) were strongly upregulated, whereas *CG7322* was modestly downregulated (Supplementary Table S4). Together, these results indicate that the *me31B^H333R^* mutation causes limited changes in whole-ovary RNA and protein abundance. Therefore, the severe fertility and germline phenotypes of the *me31B^H333R^* mutant are unlikely to result from broad disruption of the ovarian transcriptome or proteome.

### The *me31B^H333R^* mutation alters enrichment of selected Me31B-associated proteins

Because Me31B is a prominent component of germline RNP complexes and functions as an interaction hub within these assemblies [23, 27, 35, 36], we next aimed to investigate whether the *me31B^H333R^* mutation alters the composition of Me31B-associated protein assemblies. We performed immunoprecipitation followed by mass spectrometry (IP-MS) using ovarian Me31B^H333R^ and Me31B^WT^ proteins as bait. We then normalized protein abundances in the immunoprecipitates to the amount of recovered Me31B bait to enable direct comparison of co-precipitating proteins between genotypes. Differential analysis of Me31B^H333R^ and Me31B^WT^ immunoprecipitates identified a selected group of proteins with increased mutant Me31B^H333R^ association (Figure 8). Using a stringent cutoff of FDR < 0.05 and |log2FC| > 1, six proteins were significantly enriched in Me31B^H333R^ immunoprecipitates, including two established Me31B-associated protein partners, Trailer hitch (Tral) and Ypsilon schachtel (Yps) [23, 38], with log2FC values of 2.31 and 2.30, respectively (Figure 8A and Supplementary Table S5 and S6). Four ribosomal proteins, RpS26, RpS18, RpL22, and RpS6, also met these criteria, with log2FC values of 3.08, 1.62, 1.39, and 1.57, respectively (Figure 8A).

**Figure 8.**
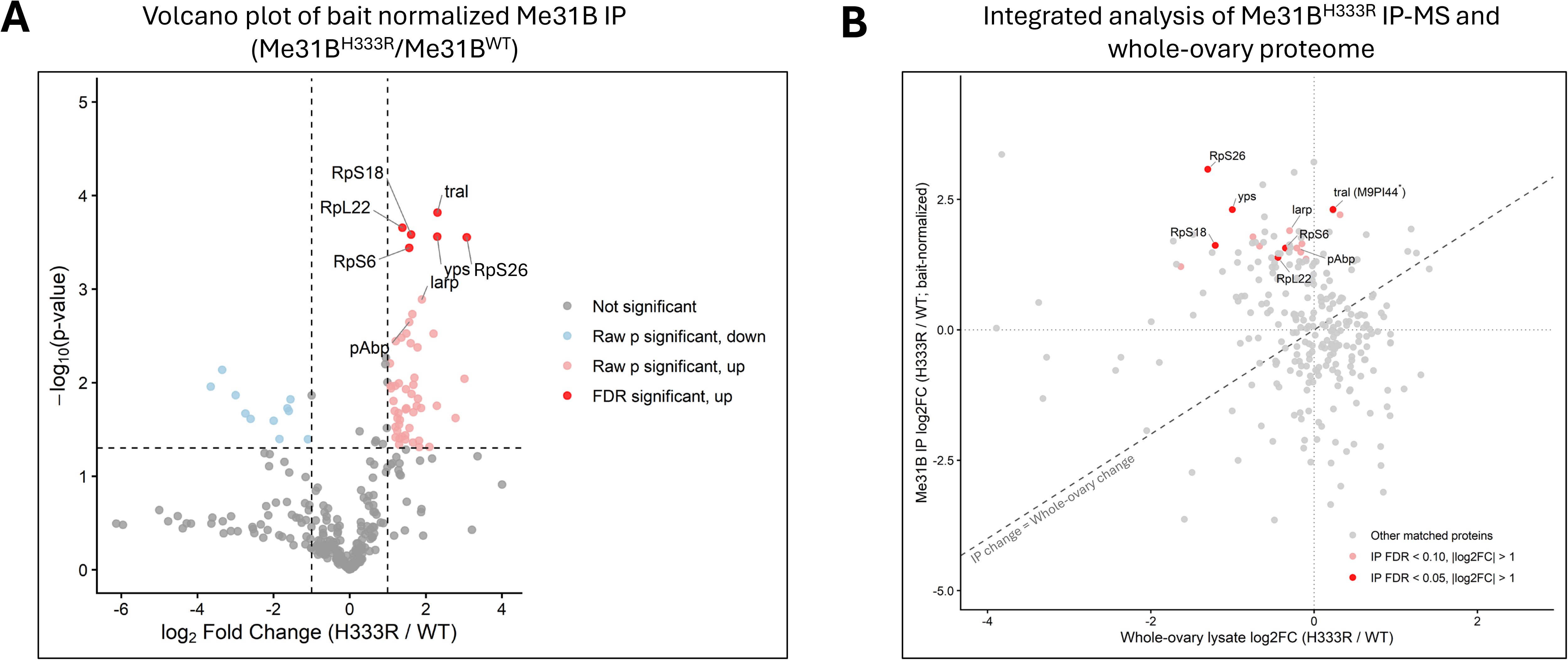
The *me31B^H333R^* mutation alters the relative abundance of select Me31B-associated proteins in bait-normalized IP-MS analysis. **(A)** Volcano plot comparing Me31B^H333R^ and Me31B^WT^ immunoprecipitates after normalization to recovered Me31B protein bait. Each point represents one quantified protein. The x-axis shows log2 fold change (Me31B^H333R^*/*Me31B^WT^), and the y-axis shows –log10 (raw p-value). The horizontal dashed line indicates the raw p-value cutoff (*p* = 0.05), and the vertical dashed lines indicate the fold-change cutoff (|log2FC| = 1). Proteins with raw p-value significance and negative log2 fold change are shown in light blue, raw p-value significant proteins with positive log2 fold change are shown in light red, and proteins meeting the FDR criterion and positive log2 fold change are shown in red. **(B)** Comparison of Me31B IP-MS enrichment with whole-ovary protein abundance changes. H333R/WT log2 fold changes from bait-normalized Me31B IP-MS were plotted against corresponding whole-ovary proteome changes. Gray points indicate other matched proteins, light red points indicate IP-MS candidates with FDR < 0.10 and |log2FC| > 1, and red points indicate those with FDR < 0.05 and |log2FC| > 1. The dashed diagonal line represents equal changes in the IP and lysate datasets. Selected candidates are labeled. For Tral (M9PI44*), the lysate value for the alternative Tral isoform M9PF14 was used as a proxy. Statistical comparison of IP and lysate changes was performed by interaction analysis as described in Materials and Methods, with results provided in Supplementary Table S6.

Using a less stringent exploratory threshold of FDR < 0.10 and |log2FC| > 1 identified additional candidates, including Poly(A)-binding protein (PABP; Log2FC = 1.57) and La-related protein (Larp; Log2FC = 1.90), as well as several additional ribosomal proteins (Figure 8A and Supplementary Table S5 and S6). Larp is physically and functionally connected with PABP in translational regulation [39–41]. The enrichment of Larp and PABP under the exploratory criteria, together with the stronger enrichment of Tral and Yps, is consistent with a selective shift in the relative composition of Me31B-associated RNP complexes and raises the possibility that the Me31B^H333R^ mutation may alter the stoichiometry of translation-related Me31B RNP assemblies.

To determine whether the IP-MS changes reflected altered protein abundance in the mutant ovary, we compared the IP-MS results with the whole-ovary proteome. Major IP-MS candidates, including Tral, Yps, Larp, PABP, and several ribosomal proteins, showed greater enrichment in Me31B^H333R^ immunoprecipitates than would be expected from their whole-ovary abundance changes (Figure 8B and Supplementary Table S6).

Of the 15 IP-MS candidates meeting the exploratory IP-MS threshold, 13 remained significantly enriched after accounting for lysate abundance differences, including the four translational regulators, Tral, Yps, Larp, and PABP (Figure 8B and Supplementary Table S6). These results support that the increased recovery of most candidate proteins in Me31B^H333R^ immunoprecipitates reflects altered association with Me31B-containing complexes rather than increased protein abundance in the mutant ovaries.

## Discussion

In this study, we generated and characterized a *Drosophila* Me31B H333R missense mutation in the conserved QxxR motif, corresponding to the human DDX6 H372R mutation associated with developmental diseases. We found that the *me31B^H333R^* mutation compromises female fertility in a dose-sensitive manner, with heterozygous females showing reduced egg hatchability and homozygous females producing eggs that fail to hatch. The homozygous phenotype is especially notable because mutant females produced externally normal eggs, indicating that an obvious defect in egg morphology cannot explain the sterility.

Importantly, this phenotype is distinct from several previously characterized *me31B* mutations that also disrupt fertility and/or embryonic development (Supplementary Table S7). Specifically, mutations affecting canonical helicase-core motifs, including *me31B^E208A^*, *me31B^DVLAAAA^*, and *me31B^R385Q^*, dominantly cause sterility with conspicuous oogenesis and embryonic defects, whereas an N-terminal deletion mutation, *me31B^C-ter^*, causes recessive sterility associated with embryonic patterning defects. In contrast, the *me31B^H333R^* mutation produces an incomplete-dominant reduction in fertility, complete sterility in homozygous females, and severe hatchability failure without obvious external egg morphology defects. Thus, the QxxR motif disruption appears to define a distinct class of Me31B functional impairment: one that strongly compromises egg competency while leaving gross egg morphology relatively intact.

Before migration to the embryonic gonad, normal germline development includes PGCs undergoing zero to two asynchronous rounds of mitosis and then becoming quiescent [42]. In examining PGCs in the embryos from *me31B^H333R^* mutant females, we found an increase in the total number of PGCs between the *me31B^H333R^*/+ mutant and the control, but no increase in the coalescing of PGCs within the embryonic gonad.

Together, this suggests that the increase in the total number of defective PGCs (cells outside the embryonic gonad) is due to an overall excess of PGCs (Figure 4). This could be due to additional PGC proliferation, which creates extra Vasa-positive cells with genetic information too diluted to navigate to the embryonic gonad. Ectopic expression of Cdc2AF (an active form of cyclin-dependent kinase, Cdc2) with either Cyclin A or Cyclin B can induce mitosis in PGCs [42], suggesting that the *me31B^H333R^*/+ mutant may affect cell cycle regulators in PGCs. However, we do not see this same observation in the *me31B^H333R^* mutant and reason that the PGCs in the *me31B^H333R^* mutant do not undergo additional proliferation, possibly due to the homozygous mutants’ more extreme effects on development in general; thus, we only see an increase in defective cells rather than an increase in the total number of cells. Regardless, our results reflect abnormal development in both the *me31B^H333R^*/+ and *me31B^H333R^* mutants.

In our smFISH experiments, we see that homotypic clustering of key germ plasm mRNAs, *nos* and *pgc,* is unaffected by the *me31B^H333R^* mutation. An estimated 59 additional types of mRNAs become localized to the germ plasm [43]. Thus, we cannot rule out potential defects in the localization and clustering of other germ plasm mRNAs in the *me31B^H333R^* mutant. In *Drosophila*, homotypic clustering and germ granule assembly rely on the mRNA localization and local posterior translation of *oskar*, the master nucleator of germ plasm formation [44]. Previous research shows that reducing Oskar levels reduces the sizes of *nos* and *pgc* homotypic clusters [32, 33]. Because *nos* and *pgc* localization and clustering were unaffected in the *me31B^H333R^* mutant, our results suggest that *me31B^H333R^* mutation does not affect the localization or translation of posteriorly localized *oskar*. Thus, we tentatively conclude that the *me31B^H333R^* mutant phenotypes are not caused by a broad effect on the germ plasm assembly process. Moreover, the preserved accumulation and posterior localization of Nos protein further support the idea that *me31B^H333R^* does not produce a broad failure in maternal mRNA translational control, at least for the *nos* regulatory program examined here. This result is notable because *nos* mRNA is regulated by Me31B-containing translational repression complexes [35], making Nos a relevant readout for assessing Me31B-dependent post-transcriptional regulation in the germ plasm. Nevertheless, this interpretation should be interpreted cautiously, since Nos is a single target and anti-Nos staining reflects steady-state protein distribution rather than direct translational activity.

At the molecular level, the *me31B^H333R^* mutation did not significantly reduce Me31B protein abundance or broadly remodel the ovarian transcriptome or proteome. RNA-seq and whole-ovary proteomics revealed limited, selective changes, with no functional category emerging from the differentially expressed transcripts and differentially abundant proteins, and only minimal overlap between them. This lack of global remodeling argues against a model in which the hatchability defect results from widespread disruption of overall maternal gene expression. Instead, these findings better fit a model in which *me31B^H333R^* disrupts more specific regulatory processes, such as post-transcriptional or translational control, that may act late in oogenesis or during early embryogenesis and may not be readily captured by steady-state, whole-ovary profiling. Notably, the only genes showing concordant upregulation at both the transcript and protein levels were *TotA* and *TotC*, two stress-response factors [37], suggesting that their induction may reflect a secondary stress-associated response. Together, these results suggest that severe reproductive phenotypes caused by *me31B^H333R^* can arise without broad transcriptome- or proteome-level disruption, emphasizing the need for temporally and mechanistically focused approaches to determine how the QxxR motif supports Me31B function during egg viability and early embryogenesis.

The Me31B IP-MS analysis provides a potential molecular explanation for how the *me31B^H333R^* mutation may impair germline development without causing broad transcriptomic or proteomic disruption. The increased enrichment of several post-transcriptional and translational regulators, including Tral, Yps, PABP, and Larp, together with multiple ribosomal proteins in Me31B^H333R^ immunoprecipitates, raises the possibility that the mutation alters the relative composition or regulatory state of Me31B-associated RNP complexes. This notion is further supported by integration of the IP-MS data with the whole-ovary proteome, which showed that the increased recovery of Tral, Yps, PABP, and Larp, as well as most other IP-MS candidates, remained significant. Me31B functions within RNA-protein assemblies that regulate maternal mRNA translation during oogenesis and early embryogenesis, and Tral and PABP are established components of Me31B-containing repressive RNPs [26, 38]. Previous studies have proposed that Me31B and Tral cooperatively associate with target RNAs and contribute to translational silencing by organizing repressive RNP structures [26, 35]. In this context, altered association of these translational regulators could potentially shift the balance between translationally repressed and translationally engaged RNP states, leading to inappropriate regulation of maternal transcripts required for oogenesis or early embryonic development. Such a mechanism could explain how the *me31B^H333R^* mutation severely compromises the egg’s developmental competence while producing only limited global molecular changes. The enrichment of ribosomal proteins in the Me31B^H333R^ immunoprecipitates should be interpreted cautiously, because ribosomal proteins are abundant and can frequently co-purify in RNA-associated immunoprecipitation experiments. Nevertheless, the coordinated enrichment of multiple ribosomal proteins (RpS18, RpS26, RpS6, RpS2, RpL24, RpL4, RpL10, RpL23A, and RpL19) under either stringent or exploratory IP-MS criteria (Supplementary Table S6) may reflect altered association of translation-related components with Me31B^H333R^-containing complexes.

## Materials and Methods

### Fly strain generation by CRISPR gene editing

Mutant *me31B^H333R^* and *me31B^WT^* control *Drosophila* strains were generated by using the CRISPR methods previously described [15]. In the constructs, a super-fold GFP (*sfGFP*) gene was positioned in-frame and downstream of the *me31B* gene so that the sfGFP protein is tagged to the C-terminus end of the expressed Me31B protein. A *DsRed* marker gene was positioned in the intergenic region downstream of the *me31B* gene. The generated strains were crossed with a 2^nd^ chromosome balancer (CyO) to establish balanced stocks. Balanced *me31B* wild-type and mutant strains were self-crossed to obtain homozygous strains.

### Female fertility assay

Fertility assays were performed as previously described [15] with minor modifications. Briefly, virgin females from *me31B^H333R^* and *me31B^WT^* control strains were collected and allowed to age for 3 – 4 days in separate vials. Each female was then put in a vial with a *w^1118^* male. The flies were transferred to a new vial every day for the next 10 days. Eggs laid and progeny hatched from each vial were counted. The hatch rate was calculated by dividing the number of progeny hatched by the number of eggs laid.

### Embryo morphology and hatchability

The embryo morphology and hatchability assays were performed as previously described [15]. Briefly, approximately 80 females from each strain were put into a small grape-agar embryo collection cage (Genesee Scientific) in the presence of 20 *w^1118^* males, and the embryos were collected on grape agar plates at 25°C after 24 hours. The embryos were counted under a dissection microscope and examined their morphology. Embryo hatchability was calculated 72 hours after laying by counting the number that developed into larvae (or later stages) or failed to develop, and subtracting the failed ones from the total number of eggs laid.

### Western blot

10 µL of ovaries from *me31B^H333R^* and *me31B^WT^* flies were homogenized in 50 µL lysis buffer (PBS, 0.1% NP-40, 1x protease inhibitor cocktail (Roche)). The lysates were centrifuged at 17,000 ×g for 15 minutes at 4°C, and 45 µL of the cleared supernatant was transferred to a new tube. Protein concentrations of the lysates were measured by using Bradford assays. The lysates were diluted to 1.5 mg/mL with lysis buffer before SDS-PAGE. The proteins were detected with rabbit anti-GFP primary antibody (Abcam) at a 1:100,000 dilution and mouse anti-rabbit HRP secondary antibody (Jackson ImmunoResearch) at a 1:10,000 dilution. Tubulin controls were detected with mouse anti-α-Tubulin (Santa Cruz) at 1:4,000 and goat anti-mouse HRP (Jackson ImmunoResearch) at 1:2,000. Imaging was performed on a Bio-Rad ChemiDoc system using chemiluminescent detection. Three biological replicates were performed. Band intensity was quantified using ImageJ (https://imagej.nih.gov/ij/) and normalized to the tubulin control.

### Embryo collection, fixation, smFISH, immunofluorescence experiments, and image quantification

For smFISH experiments, flies were first fed yeast paste for 24 hours, followed by a 1-hour embryo collection on fresh grape juice agar plates. For immunofluorescence (IF) experiments, 24-hour embryos were collected. For both sets of experiments, embryos were dechorionated, devitellinized, fixed in 4% paraformaldehyde, and stored in 100% methanol following standard established protocols [28]. Previously described smFISH probes for nos (ATTO 647N dye) and pgc (Quasar 570 dye) were used [28]. Previously established smFISH protocols were carried out as described [33, 45]. IF experiments were performed using previously described protocols [46, 47] using anti-Vasa (Boster Bio, cat #DZ41154) and secondary Alexa Fluor 568 (anti-rabbit, ThermoFisher, cat #A10042) to mark PGCs [28]. Vasa is an established PGC marker [48]. IF and smFISH samples were mounted in ProLong Glass (Life Technologies), and samples were cured for three days before being imaged. Confocal microscopy was performed using a Leica STELLARIS 5 white light laser system in photon counting mode for smFISH as described in detail [28, 32].

Generation of germ granule censuses was carried out using established image quantification techniques [31–33]. In summary, a custom MATLAB (Mathworks) program 1) identifies and quantifies the average intensity of unlocalized single transcripts, 2) homotypic clusters in the germ plasm are identified using a user-defined polygon that marks the entire germ plasm within a 15-slice z-stack (∼5 µm), and their intensities are normalized by using the average intensity of the single transcripts. The Granule Census identifies clusters that reside in the same granules by calculating the frequency that two clusters’ centroids are found within a previously established distance threshold [28, 31, 32]. Confocal images in Figure 6 were filtered using a balanced circular difference of Gaussian with a surround size of 2.2 pixels and center radius size of 1.2 pixels [31].

Nos immunostaining was performed on 2-hour embryos according to the *Drosophila* embryos IF protocols described above. Anti-Nos antibody was used at 1:2,000 dilution. Images were captured using an Olympus FV3000 confocal laser-scanning microscope using identical acquisition settings. For each embryo, a region encompassing the Nos immunofluorescence signal in posterior germ plasm was defined and measured as integrated density (IntDen) in ImageJ. Replicate measurements were averaged to obtain a single value per embryo. Values were normalized to the mean of the *me31B^WT^* control, which was defined as 100%. Genotypes were compared using a two-sided Welch’s t-test. The box-and-scatter plot was generated in R using the ggplot2 package.

### RNA sequencing and data analysis

Ovaries were dissected from the *me31B^H333R^* mutant and *me31B^WT^* control strains and flash-frozen in liquid nitrogen. RNA extraction, library preparation, sequencing, and initial data processing were performed by Azenta Life Sciences. Total RNA was extracted using the RNeasy Plus Universal Mini Kit (Qiagen), and poly(A)+ mRNA was enriched using oligo(dT) beads. Sequencing libraries were prepared using the NEBNext Ultra II RNA Library Prep Kit for Illumina and NEBNext Poly(A) mRNA Magnetic Isolation Module (New England Biolabs) and sequenced as 150-bp paired-end reads on an Illumina NovaSeq X Plus instrument. Raw reads were trimmed using Trimmomatic v0.36 and aligned to the *Drosophila melanogaster* BDGP6/dm6 reference genome using STAR v2.5.2b. Gene-level read counts were generated using featureCounts from the Subread package v1.5.2, and differential expression analysis was performed using DESeq2. Differentially expressed genes were defined as those with an adjusted *p* < 0.05 and an absolute log2 fold change > 1.

Downstream analyses and visualization were performed in R. For principal component analysis, genes with zero counts across all six samples were removed, raw counts were variance stabilized using DESeq2, and PCA was performed using the 500 genes with the highest variance. Volcano plots were generated from the differential expression results using ggplot2. Gene Ontology enrichment analysis was performed using the clusterProfiler package, with upregulated and downregulated genes analyzed separately for Biological Process and Molecular Function enrichment. All genes detected in the RNA-seq dataset were used as the background. FlyBase gene annotations were obtained using org.Dm.eg.db, and enrichment p-values were adjusted for multiple testing using the Benjamini-Hochberg method.

### Whole ovary lysate preparation and immunoprecipitation for mass spectrometry

*Drosophila* ovaries (20 µl) were dissected from the *me31B^H333R^* mutant strain and homogenized in 200 µL lysis buffer (0.1% NP-40 and protease inhibitor in phosphate-buffered saline). The lysate was clarified by centrifugation, and protein concentration was determined using the Bradford Assay. A portion of the crude whole-ovary lysate was submitted for mass spectrometry analysis. The same procedure was performed with the *me31B^WT^* control strain. For immunoprecipitation, the remaining lysate was diluted to 1.0 mg/ml with the lysis buffer and incubated with 25 µL anti-GFP agarose beads (MBL International) for 120 minutes at 4°C on a rotary shaker. At the end of the incubation, the beads were washed twice with lysis buffer supplemented with 150 mM NaCl and once with PBS. The beads were then subjected to on-bead digestion followed by mass spectrometry.

### Protein processing, mass spectrometry, and database search

Samples were resuspended and denatured in 8 M urea with 100 mM ammonium bicarbonate (pH 7.8). Disulfide bonds were reduced by incubation for 45 min at 57°C with a final concentration of 10 mM Tris (2-carboxyethyl) phosphine hydrochloride (Catalog no C4706, Sigma-Aldrich). A final concentration of 20 mM iodoacetamide (Catalog no I6125, Sigma-Aldrich) was then added to alkylate the side chains, and the reaction was allowed to proceed for one hour in the dark at 21°C. Samples were diluted to 1 M urea in 100 mM ammonium bicarbonate (pH 7.8). Trypsin (V5113, Promega) was added, and the samples were digested for 14 hours at 37°C. For the on-bead digestions, a total of 0.5 µg trypsin was used, while for the lysate samples, 2.0 µg was used.

Individual samples were desalted using ZipTip pipette tips (EMD Millipore), dried down, and resuspended in 0.1% formic acid. Fractions were analyzed by LC-MS on an Orbitrap Fusion Lumos equipped with an Easy NanoLC1200 HPLC (ThermoFisher Scientific). Buffer A was 0.1% formic acid in water. Buffer B was 0.1% formic acid in 80% acetonitrile. For the immunoprecipitation samples, peptides were separated on a 30-minute gradient from 0% B to 3% B. Precursor ions were measured in the Orbitrap with a resolution of 120,000. For the lysate samples, this gradient went for 90 minutes.

Fragment ions were measured in the Orbitrap with a resolution of 15,000. The spray voltage was set at 1.8 kV. Orbitrap MS1 spectra (AGC 1×10^6^) were acquired from 350-2000 m/z, followed by data-dependent HCD MS/MS (collision energy 30%, isolation window of 2 Da) for a three-second cycle time. Charge state screening was enabled to reject unassigned and singly charged ions. A dynamic exclusion time of 30 seconds was used to discriminate against previously selected ions.

The LC-MS/MS data were searched against the Uniprot *Drosophila melanogaster* proteome (downloaded 07/2020) using Proteome Discoverer (v2.5.0.400). Trypsin was set as the protease specificity, allowing for two missed cleavages.

Carbamidomethylation of cysteine residues was set as a static modification. Oxidation of methionine, pyroglutamine formation on peptide-terminal glutamine, acetylation of the protein amino terminus, and loss of the protein amino-terminal methionine residue were set as dynamic modifications. The precursor mass tolerance was 10 ppm, and the fragment ion tolerance was 0.04 Da. Error rates were determined using Percolator with an FDR rate of 0.01.

### Integrated analysis of Me31B IP-MS and whole-ovary proteomics

Bait-normalized IP-MS data were integrated with the whole-ovary proteomics dataset in R to distinguish changes in Me31B-associated protein enrichment from changes in total ovarian protein abundance. Proteins were matched between datasets by protein accession, and H333R/WT changes in the IP and lysate datasets were compared statistically to identify proteins whose IP enrichment exceeded changes in total ovarian abundance. The interaction p-values from this comparison were adjusted using the Benjamini–Hochberg method, with FDR < 0.05 considered significant. For visualization, IP log2 fold changes were plotted against corresponding whole-ovary lysate log2 fold changes. Volcano and scatter plots were generated in R using the ggplot2 package. For Tral accession M9PI44, which lacked an exact lysate accession match, the alternative Tral isoform M9PF14 was used as a proxy and annotated accordingly.

### Use of AI-Assisted Tools

The authors used an AI-assisted tool, ChatGPT (OpenAI), to assist with language editing and R scripting for data visualization from author-provided datasets. The tool was not used to generate experimental data, conduct analyses independently, or independently determine the study’s biological conclusions. The authors reviewed, verified, and approved all data inputs, R code, analysis parameters, statistical results, biological interpretations, figures, and tables.

## Author Contributions

MG conceived and directed the project. MG and MGN wrote the manuscript with input from all authors. RM, ASM, and ANT performed the fertility assays. RM performed the RNA-seq sample preparation. AM, CD, and JE generated the CRISPR strains. JCT, MG, DG, and BN performed mass spec experiments and analyzed the data. RM, YM, and BP performed the Western blot experiments. EK, AF, IF, and AI performed the hatchability assays. MGN, MMM, and ALS performed PGC microscopic analysis and smFISH experiments. RM, ASM, AAM, AYK, and MG performed the Nos microscopic experiments and analyzed the data. All authors discussed the results and reviewed the manuscript.

## Data Availability

Strains and plasmids are available upon request. The authors affirm that all data necessary for confirming the conclusions of the article are present within the article, figures, tables, supplementary materials, or deposited in data repositories. RNA-seq data generated in this study have been deposited in the NCBI Gene Expression Omnibus (GEO) under accession GSE343730.

## Acknowledgements

We thank members of the Gao lab for helpful discussions and feedback during the development of this project. We thank current and former members of the Gao laboratory for their support and assistance throughout this project. We are grateful to Jenna Ibrahim for technical assistance and contributions to routine laboratory operations that supported the completion of this work. We thank the Center for Biological Imaging at Kean University for assisting with image acquisition and all the members of the Niepielko Lab for their helpful comments and fruitful discussions. We thank Dr. Ruth Lehmann lab for the kind gift of the Nos antibody.

Institutes of Health AREA grant (2R15HD092925-02A1) to MG and MGN and institutional support from Kean University and Indiana University Northwest. Support for The Niepielko Lab is also provided by CAREER Award 2237390 from the National Science Foundation.

Any opinions, findings, and conclusions expressed in this material are those of the author and do not necessarily reflect the views of the funding agencies.

